# Convergent reduction in skeletal density during benthic to pelagic transitions in Baikal sculpins

**DOI:** 10.64898/2026.01.22.701097

**Authors:** Brayan A. Gutierrez, Olivier Larouche, Sara Loetzerich, Mackenzie E. Gerringer, Kory M. Evans, Andres Aguilar, Sergei Kirilchik, Michael W. Sandel, Jacob M. Daane

## Abstract

Habitat transitions are a major driver of morphological evolution. Teleost fishes have repeatedly transitioned from benthic to pelagic habitats, often evolving predictable changes in body shape that enhance hydrodynamic efficiency. While freshwater sculpins (Cottidae, Perciformes) are usually benthic, two genera in Lake Baikal, *Comephorus* and *Cottocomephorus*, have independently evolved into midwater niches. As sculpins lack a swim bladder, these lineages instead improved buoyancy through reduced skeletal density and increased lipid stores. Using micro-computed tomography and two-dimensional morphometrics, we characterized skeletal evolution across the Baikal sculpin radiation. We found that parallel changes in bone mineral density and microstructure independently evolved in the two pelagic clades. Density reductions occurred throughout the skull in pelagic species. The basibranchials and neurocranium exhibited the lowest overall bone density across all cranial elements. While the jaws maintained the highest absolute density values among the bones we measured, they also showed the greatest proportional reduction in density associated with pelagic habitat use, with a 56.86% decrease in percentage hydroxyapatite and a 21.39% increase in porosity. Morphometric analyses further identified convergence toward an elongate body shape, reduced and posteriorly shifted eyes, and elevated fin insertion in pelagic taxa. These results demonstrate a repeated skeletal lightening and body shape changes accompanying benthic-to-pelagic transitions. This pattern mirrors other benthic-to-pelagic transitions in teleosts that lack swim bladders, highlighting shared biomechanical and microstructural solutions to life in the open water.

## Introduction

The skeleton serves as a critical scaffold for vertebrate morphology, providing structural support, protection for internal organs, anchorage points for muscles, and a reservoir for key minerals such as calcium and phosphorus (Šromová, Sobola, & Kaspar, 2023). Because of these essential functions, skeletal elements could be expected to experience stabilizing selection, leading to structural conservatism across evolutionary time (Sansalone et al., 2022). However, many major evolutionary innovations and ecological transitions in vertebrates have arisen through modifications of skeletal structures, allowing lineages to diversify, adapt to new environments, and evolve novel morphologies (Dial, Shubin, & Brainerd, 2015).

Teleost fishes, which account for nearly half of all extant vertebrate species (Nelson, Grande, & Wilson, 2016), exhibit striking diversity in skeletal form and function. The skeleton typically comprises only three to five percent of body mass in bony fishes, but bone is far denser and heavier than other tissues and generates negative buoyancy (Reynolds & Karlotski, 1977). Buoyancy has a major impact on the ability of individuals to station hold, swim horizontally, and stabilize posture (Di Santo & Goerig, 2025; Gleiss, Potvin, & Goldbogen, 2017; Sato, Aoki, Watanabe, & Miller, 2013; P. W. Webb, 2005; Weihs, 1973). Thus, density has a pervasive influence on aquatic locomotion, driving adaptations in both morphology and behavior (Pelster, 2009).

To counteract the weight of dense tissues, most teleosts use inflatable swim bladders to maintain neutral buoyancy (Pelster, 2009). However, the swim bladder has been lost independently at least 30 times in teleost evolution (McCune & Carlson, 2004), often in benthic species that rest on the substrate or in deep-sea fishes where the high pressures make gas inflation difficult (Priede, 2018; Scholander & Van Dam, 1954). In these cases, alternative buoyancy-control mechanisms have evolved (Pelster, 2009). One such buoyancy control mechanism is the reduction of skeletal density, a buoyancy adaptation also found in non-teleost vertebrate taxa, including deepwater cetaceans (Gray, Kainec, Madar, Tomko, & Wolfe, 2007; Pelster, 2009).

Changes in skeletal density are common among fishes and can take multiple forms including reductions in bone size and thickness, increased porosity, or lowered mineral content (Eastman, Witmer, Ridgely, & Kuhn, 2014; M. E. Gerringer et al., 2021; Martin, Dias, Summers, & Gerringer, 2022). Further, fish bones vary widely in structure, appearing spongy, compact, intermediate between bone and cartilage, or consisting primarily of connective tissue or persistent hyaline cartilage rather than fully ossified tissue (Eastman et al., 2014; Meunier & Huysseune, 1991). Changes in structure and overall density are often not uniform across the skeleton, reflecting structure-specific trade-offs between minimizing weight and preserving essential functions such as protection and muscle attachment.

Adaptive changes in skeletal density have been drivers of diversification. The cryonotothenioid, or Antarctic notothenioid (Notothenioidei, Perciformes), radiation in the Southern Ocean exemplifies this pattern. A central axis of cryonotothenioid diversification involved vertical habitat expansion to exploit pelagic prey (Eastman, 1993). Notothenioids lack swim bladders, instead achieving buoyancy through reduced skeletal density and increased lipid accumulation (Eastman, 2024; Eastman et al., 2014). Notably, initial reductions in skeletal density evolved prior to the adaptive radiation in the Antarctic Southern Ocean, which is thought to have facilitated subsequent diversification into pelagic niches (Daane et al., 2019).

A similar, though less studied, adaptive radiation has occurred among the sculpins (Cottidae, Perciformes) in Lake Baikal. At 1,642 meters (m) maximum depth (De Batist, Canals, Sherstyankin, Alekseev, & INTAS Project 99-1669 Team, 2006), Lake Baikal is the world’s oldest and deepest freshwater lake. Unique among lake habitats, Lake Baikal is well-oxygenated along the entire water column. This oxygenation supports life in marine-like habitats such as hydrothermal vents, cold seeps, and bathypelagic zones, and has fostered more than 1,500 endemic species and multiple animal adaptive radiations (Macdonald, Yampolsky, & Duffy, 2005; Ravens, Kocsis, Wüest, & Granin, 2000; Sandel et al., 2025; Stelbrink et al., 2015). Cottid sculpins dominate the fish fauna of Lake Baikal, comprising 57% of fish species diversity (Matveyev & Samusenok, 2015). Like notothenioids, the Baikal sculpin radiation is marked by vertical habitat expansions, particularly colonization of deep benthic habitats, with elevated speciation rates in the bathybenthic clade (Sandel et al., 2025).

Sculpins have also diversified into the vast pelagic zone of Lake Baikal, where 80% of the lake’s surface area is at depths exceeding 300 m (Kolokoltzeva, 1961; Valentina G. Sideleva, 2003) (**Fig. 1A**). In this habitat, six pelagic sculpin species dominate, comprising ∼80% of total fish biomass (Valentina G. Sideleva, 2003). The six pelagic sculpins are distributed between two genera, *Comephorus* and *Cottocomephorus*, and represent two independent benthic-to-pelagic transitions (Sandel et al., 2025)*. Comephorus* species are fully pelagic, engage in diel vertical migrations for nocturnal feeding, and have evolved ovoviviparity, which removes a dependency on benthic egg nesting (Valentina G. Sideleva, 2003; Taliev, 1955). In contrast, *Cottocomephorus* species feed in midwater during the day but remain tied to the benthos for spawning, nesting, and resting (Valentina G. Sideleva, 2003). The two genera also differ in their primary pelagic prey items, which is reflected in morphological traits relevant for trophic ecology. As adults, *Comephorus* preys on a combination of large and mobile *Macrohectopus* amphipods and other fish (Anoshko, Dzyuba, & Melnik, 2002). To facilitate prey capture, *Comephorus* has a large mouth opening with elongated jaws (16% of body length) that are lined on the exterior with needle-like teeth (Aleev, 1963; Valentina G. Sideleva, 2003). *Cottocomephorus* largely target zooplankton and *Epischurella* copepods, possessing a mouth half the size of that in *Comephorus*, toothless outer jaws, and relatively long gill rakers (Valentina G. Sideleva, 2003; V. G. Sideleva & Kozlova, 1989; Taliev, 1955).

**Fig. 1.**
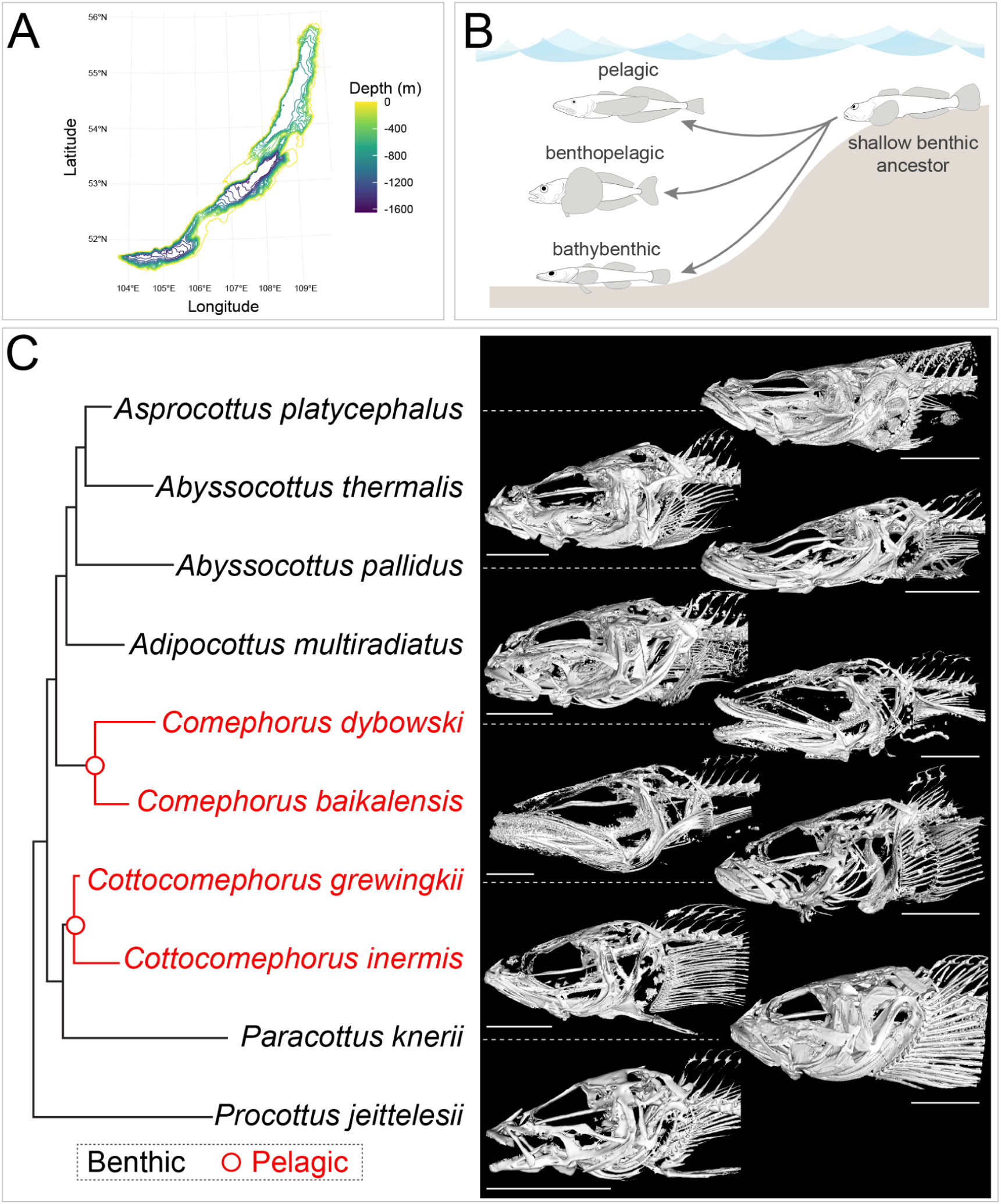
Benthic-to-pelagic transitions during the Baikal sculpin adaptive radiation. A) Bathymetric map of Lake Baikal [data from (De Batist et al., 2006)]. B) The adaptive radiation of the Baikal sculpins from a shallow benthic common ancestor into bathybenthic, benthopelagic, and pelagic species. C) Cladogram and skull reconstructions at optimum brightness for each scan from selected species of Baikal sculpins. Red indicates pelagic (*Comephorus*) or benthopelagic (*Cottocomephorus*) species, black indicates benthic species. Cladogram adapted from (Sandel et al., 2025). Skeleton brightness optimized for each scan. Scale bars in C represent 10 mm.

Like notothenioids, pelagic Baikal sculpins diversified from a negatively buoyant benthic common ancestor lacking a swim bladder. Whereas benthic sculpins typically have skeletal ash weights above 3%, pelagic forms fall below this threshold, with *Comephorus baikalensis* achieving near-neutral buoyancy with skeletal ash weight as low as 1.6-2.5% and a specific gravity of 1.010 (Valentina G. Sideleva, 2003; Taliev, 1955). *Comephorus* is reported to have thinner neurocranial and pectoral fin bones and increased bone porosity, though details are scarce (Valentina G. Sideleva, 2003).

Here, we characterize skeletal density variation of Baikal sculpins to assess phylogenetic patterns, trait-habitat associations during adaptive radiation into the pelagic environment, and underlying modes of density reduction. This study provides new insights into how skeletal shifts facilitate benthic-to-pelagic transitions, with broader implications for understanding adaptive radiations in aquatic systems.

## Materials and Methods

### Species sampling and habitat scoring

Baikal sculpin specimens were collected from Lake Baikal via trawling, snorkeling, SCUBA, dipnets, funnel traps, and seines, fixed in 10% formalin and stored in 70% ethanol as described previously (Sandel et al., 2025; St. John et al., 2022). We analyzed replicates from ten species in Lake Baikal from four out of the five taxonomic tribes (Bogdanov, 2023): Comephorini (*Comephorus baikalensis* [n=3 individuals], *Comephorus dybowski* [n=3]), Abyssocottini (*Adipocottus multiradiatus* [n=4]*, Asprocottus platycephalus* [n=3], *Abyssocottus pallidus* [n=3], *Abyssocottus thermalis* [n=2]), Cottocomephorini (*Cottocomephorus grewingkii* [n=6], *Cottocomephorus inermis* [n=4]) and Cottini (*Procottus jeittelesii* [n=4]*, Paracottus knerii* [n=5]).

Species were classified as either benthic (n = 6 species) or pelagic (n = 4) (**Fig. 1B, C**) (Valentina G. Sideleva, 2003). Within the pelagic group, species of the genus *Comephorus* represent a true transition to a pelagic lifestyle, while species of the genus *Cottocomephorus* are considered benthopelagic, occupying both benthic (bottom-dwelling) and pelagic (open water) zones (Valentina G. Sideleva, 2003). Because these latter species are active swimmers that hover above the substrate, we expect they should experience similar selective pressures related to buoyancy as the pelagic species.

### Scanning of specimens

We performed micro-computed tomography (micro-CT) scans of selected Baikal sculpin species using a Bruker SkyScan 1273 micro-CT scanner configured with 70 kV X-ray voltage, 214 µA current, medium focal spot size, and 0.3° rotation step. Scan lengths ranged from 145 to 245 millimeters (mm) with voxel sizes between 12 and 20 micrometers (µm), optimized for each specimen’s size. Two calcium hydroxyapatite phantom markers (25% and 75% density) were included in each scan to calibrate bone density measurements to scan brightness. Micro-CT data were reconstructed using Bruker NRecon v.2.1.0.1, cropping the region from snout to the sixth or seventh vertebra. Beam hardening correction (44%) minimized polychromatic X-ray artifacts, with dynamic image range set between 0.00 and 0.095 for optimal contrast. All image processing and segmentation used NRecon, DataViewer, and Amira Version 6.0.1 on Windows OS, adapting methodologies from Martin et al. (2022) and Buser et al. (2020) for compatibility with current Bruker software. For additional details on scanning and image processing, see **Supplementary Methods**.

### Bone Mineral Density, Bone Volume Fraction, and Bone Thickness

We estimated three metrics of skeletal density and morphology: bone mineral density, porosity, and thickness. We focused on cranial elements critical for feeding and structural integrity, including the preopercle, ceratohyal, fifth ceratobranchial, dentary, dorsal neurocranium, basibranchials, and cleithrum (see **Fig. 2A**). The dorsal neurocranium makes up the rigid roof of the skull that protects the brain. The dentary is the main, tooth-bearing bone of the lower jaw. The preopercle supports the posterior of the cheek just ahead of the gill cover. The cleithrum is a major bone of the pectoral girdle that connects the pectoral fins to the back of the skull through intervening elements. The ceratohyal, a key component of the hyoid arch, forms part of the mouth floor and connects ventrally to the basibranchials, a series of midline bones that support and link the gill arches. Finally, the fifth ceratobranchial is a modified, often tooth-bearing bone that serves as a pharyngeal jaw for processing food.

**Fig. 2.**
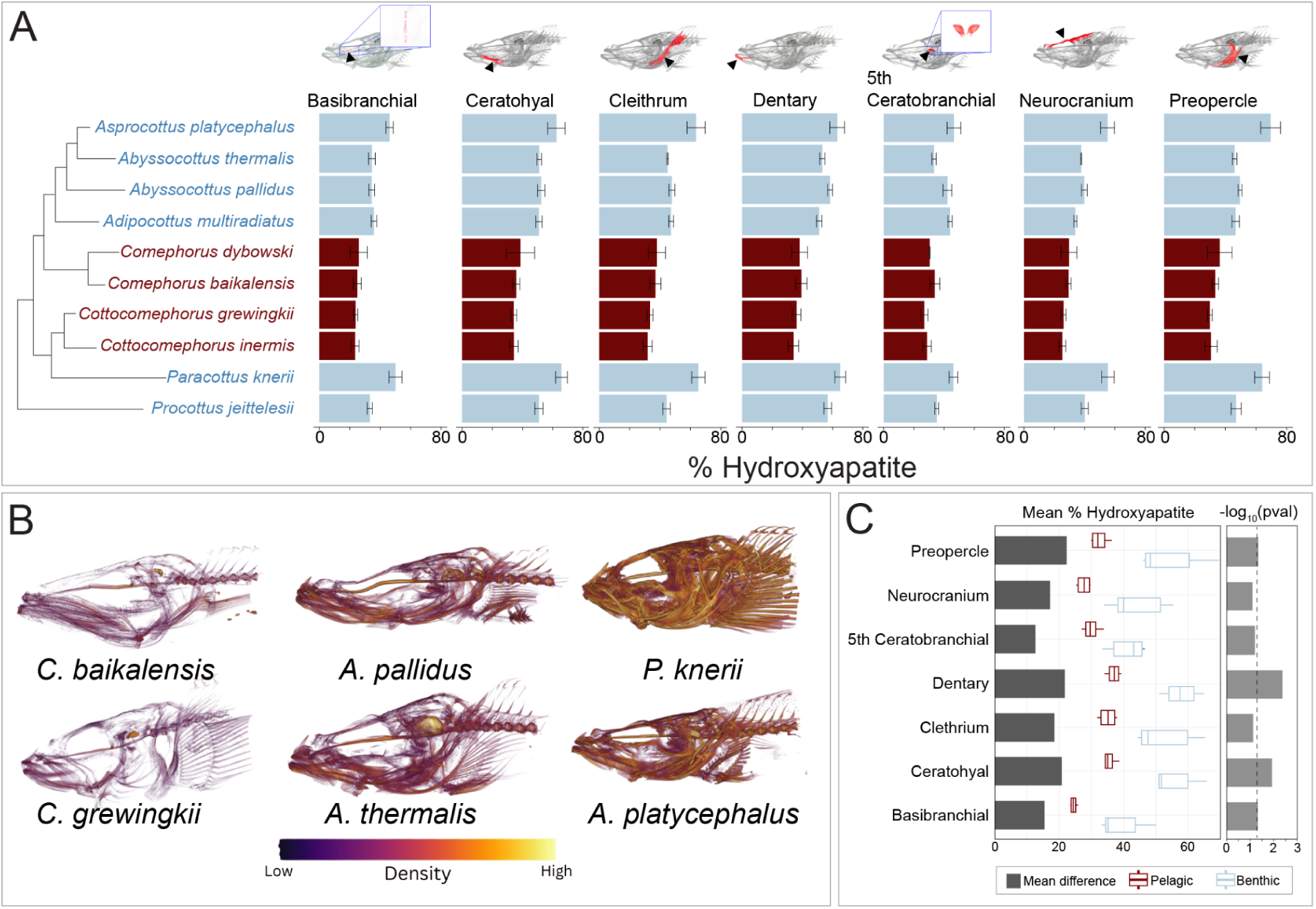
Patterns of skull bone mineral density across the Baikal sculpin radiation. A) Mean percentage of hydroxyapatite, comparing benthic (blue) and pelagic (red) groups. B) Skull density heatmaps of representative Baikal sculpins, including two pelagic species (*C. baikalensis* and *C. grewingkii*) and four benthic species (*A. pallidus*, *A. thermalis*, *A. platycephalus*, and *P. knerii*). C) Mean differences in hydroxyapatite percentage between benthic and pelagic groups, with ANOVA *P*-values transformed via −log₁₀ for visualization (**Table S2**). Dashed line indicates a *P*-value of 0.05.

Bone mineral density was assessed using the hydroxyapatite phantoms as reference standards, converting grayscale intensity values from the scan data into mineral density measurements through relative pixel brightness comparisons, which translates to a percentage of hydroxyapatite value (% HA). For each scan, we determined mean pixel brightness (MPB) values from these markers in full and developed scan-specific linear models using mean pixel brightness as the predictor and known % HA as the response variable. These models allowed us to convert mean pixel brightness measurements from segmented bones into predicted % HA values, providing standardized mineral density assessments across skeletal elements (Martin et al., 2022).

Bone volume fraction (BVF) offers valuable insight into bone density by quantifying the spatial organization of bone tissue. We used this metric to assess and compare bone porosity across Baikal sculpins. Using 3D Slicer (Version 5.6.2) (Kikinis, Pieper, & Vosburgh, 2014), we obtained the two necessary measurements for BVF calculations: bone volume (BV) and total volume (TV). BV represents the actual segmented bone volume, where higher porosity results in lower BV values. TV measures the same region with all voids filled, representing a hypothetical solid bone structure (**Fig. S1**). The BVF is derived from the BV/TV ratio, ranging from near-zero (highly porous) to one (completely solid). We computed BVF for four of our seven target bones, excluding the cleithrum and preopercle due to issues in void filling accuracy for arched bones and the basibranchial due to its small size and low mineral density.

We conducted comparative thickness analyses of the same four bones examined in our BVF assessment across Baikal sculpin species. For each segmented bone, thickness values were computed in 3D Slicer and stored as a scalar field on the surface model, with one thickness value assigned to each mesh vertex. Using a custom Python script, we extracted these per-vertex thickness scalars and calculated the mean thickness across all surface points for each bone. These values represent average geometric bone thickness (in millimeters) derived directly from the model geometry rather than material density. The resulting mean thickness values provide a standardized, specimen-level descriptor of bone geometry suitable for cross-taxon comparisons and correlation with other structural metrics such as bone volume fraction.

Bone mineral density was computed using Amira (Version 2020.3) on Windows OS. We used built-in segmentation, thresholding, and material statistics tools to help calculate mean pixel brightness and use that metric to translate it to our % HA measurements. Our BVF and thickness metrics were computed using 3D Slicer on Windows OS. We utilized built-in modules including Volume Rendering, Models, Data, Segment Editor, Segment Statistics, Simple Filters, and Probe Volume With Model. For enhanced segmentation and analysis capabilities, we additionally installed the Slicer Morph and Surface Wrap Solidify modules. For further details on all density calculations, see **Supplementary Methods**.

### Geometric morphometrics analysis

We used two-dimensional geometric morphometrics to examine overall body shape differences between benthic and pelagic sculpins and to investigate the potential relationship between body size and bone properties.

All specimens were photographed in lateral view following a standardized protocol. We positioned each specimen on a dissecting tray using insect pins to stabilize them and mark anatomical landmarks that may be difficult to visualize in photographs (e.g., the base of median fins that often could not be fanned without risking damage to the preserved specimens). Each photograph included a ruler for scale calibration. Images were captured using an Olympus OM D E M1 Mark II camera equipped with an OM System M.Zuiko Digital ED 25mm F1.8 II lens mounted on a copy stand. To maintain optimal image quality across different sized specimens, we adjusted the camera height to 37.5 cm for small specimens, 43.2 cm for mid-sized specimens, and 48.3 cm for large specimens, ensuring consistent focus and framing throughout our dataset.

We digitized morphological features using tpsDig2 (v 2.32) (Rohlf, 2017), using 15 fixed landmarks and 53 sliding semilandmarks to comprehensively capture body shape characteristics and fin positioning across all specimens (**Fig. S2**, **Table S1**). To standardize our shape data, we performed a generalized Procrustes analysis that accounted for variations in specimen size, position, and orientation, resulting in aligned shape coordinates and centroid size measurements for each individual. Subsequently, we computed species-level average shapes using the mshape function in the geomorph package (Adams & Otárola-Castillo, 2013).

The following replicates were used per species for geometric morphometrics: *C. baikalensis* (n=5 individuals), *C. dybowski* (n=5), *A. multiradiatus* (n=7)*, A. platycephalus* (n=9), *A. pallidus* (n=6), *A. thermalis* (n=2), *C. grewingkii* (n=8), *C. inermis* (n=4), *P. jeittelesii* (n=4)*, and P. knerii* (n=7).

### Comparative methods

We conducted all comparative analyses using a recent phylogeny (Sandel et al., 2025), which we pruned to include only the species present in each dataset. To examine habitat effects on bone metrics, we performed one-way phylogenetic ANOVAs using the aov.phylo function from the geiger package (v 2.0.11) in R (Pennell et al., 2014). We assessed statistical significance by comparing observed values against a null distribution generated from 1000 simulations under a Brownian motion model, running 1000 different ANOVA models to obtain an average *P*-value. These analyses compared average bone densities between benthic and pelagic Baikal sculpin species. Prior to ANOVA, normality of each response variable was assessed using Shapiro–Wilk tests conducted separately for each bone, consistent with the bone-specific ANOVA design. All measurements showed no significant departure from normality (*P* > 0.05) except for neurocranium porosity (*P* = 0.04505). We applied identical approaches to evaluate habitat effects on average bone porosity and thickness, creating separate ANOVA models for each bone of interest.

To investigate relationships between bone properties, we conducted phylogenetic multiple linear regressions using the phylolm function from the phylolm R package (Ho & Ané, 2014). These analyses tested for linear relationships between bone density (response variable) and both porosity and thickness (predictor variables), incorporating data from all four bones measured for porosity and thickness. This allowed us to assess whether bone perforation patterns and thickness influence mineral density.

We also examined potential correlations with body size by calculating Spearman correlation coefficients between each bone metric and centroid size derived from our 2D geometric morphometrics analysis.

We analyzed shape variation patterns in Baikal sculpins using two complementary approaches. First, we conducted a principal components analysis of shape coordinates using the gm.prcomp function in geomorph to identify major axes of morphological variation. Second, we performed phylogenetic Procrustes ANOVAs with the procD.pgls function (geomorph package) to test for habitat effects on body shape, assessing significance against a null distribution generated from 1000 permutations of tip data. The PCA results revealed that PC1 effectively distinguished benthic and pelagic species, prompting additional phylogenetic ANOVAs in geiger using species scores along this first principal component to further quantify habitat related shape differences.

## Results

### Independent reductions in skeletal density across pelagic sculpins

We used micro-CT scans to generate 3D skeletal reconstructions from multiple individuals of 10 species spanning the Baikal sculpin adaptive radiation (**Fig. 1C**). From these reconstructions, we measured craniofacial skeletal density using three primary metrics: mineral density, porosity, and thickness. These patterns were then compared across the Baikal sculpin radiation to identify phylogenetic trends in skeletal density.

Mineral density was estimated relative to calcium hydroxyapatite phantom markers. Mineral density varied substantially across the dataset, both among individual bones and between species. The basibranchials and neurocranium consistently had the lowest densities, while the dentary exhibited the highest mineral density values (**Fig. 2**). There were distinct species-specific differences in overall mineral density that were consistent across all measured bones. *P. knerii* and *A. platycephalus* showed the highest mineral density, while *A. pallidus* and *A. thermalis* had intermediate levels (**Fig. 2A,B**). The pelagic species in the dataset (*C. grewingkii*, *C. inermis*, *C. dybowski*, and *C. baikalensis*) exhibited the lowest mineral densities for each bone (**Fig. 2A,C**). Although mineral density was universally reduced in pelagic sculpins compared to benthic taxa, certain elements showed disproportionately larger decreases compared to benthic cottids, particularly the preopercle, dentary, and ceratohyal (**Fig. 2C**). These reductions in mineral density are convergent between *Comephorus* and *Cottocomephorus* (**Fig. 2A**).

We estimated bone porosity using bone volume fraction, defined as the proportion of a segmented skeletal element’s volume that is occupied by mineralized tissue. Bone volume fraction values approaching one indicate low porosity, while values closer to zero reflect increasing porosity. The dentary, ceratohyal, and fifth ceratobranchial maintained relatively consistent porosity levels with minimal variation between species (**Fig. 3**). The neurocranium displayed the greatest variation in porosity, with increases in the pelagic sculpins (*C. baikalensis*, *C. dybowski*, *C. grewingkii*, and *C. inermis*) as well as the benthic *A. multiradiatus* (**Fig. 3A**). The dentary also showed distinct porosity differences, though not clearly delineating between benthic or pelagic habitats (**Fig. 3A,C**). As examples of habitat-specific patterns observed across the various bones, benthic *A. pallidus* had among the lowest porosity, benthopelagic *C. grewingkii* showed an intermediate porosity that is common among other benthic sculpins, while *C. baikalensis*, the only sculpin near neutral buoyancy (Valentina G. Sideleva, 2003; Taliev, 1955), exhibited the highest porosity, which was consistent across multiple bones (ceratohyal, dentary, neurocranium; **Fig. 3A,B**).

**Fig. 3.**
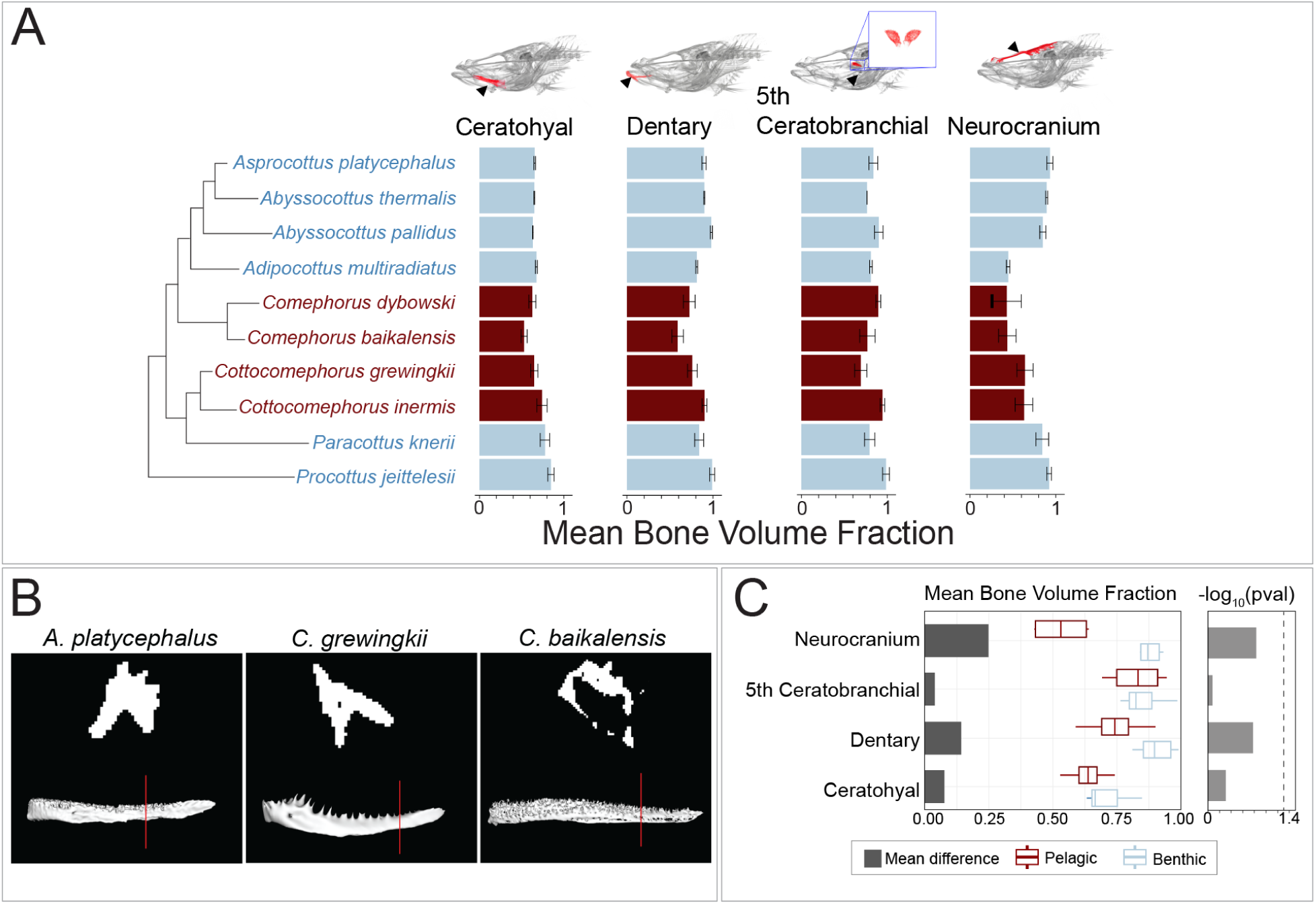
Patterns of bone porosity across the Baikal sculpin radiation. A) Mean bone volume fraction, a measure of porosity, in selected Baikal sculpin species, comparing benthic (blue) and pelagic (red) groups. B) Dentary cross-sections (taken at the plane indicated by red lines) highlighting differences in porosity between representative species. C) Mean BVF differences between benthic and pelagic groups, with ANOVA *P*-values transformed via −log₁₀ for visualization (**Table S2**). Dashed line indicates a *P*-value of 0.05.

Bone thickness also varied across the Baikal radiation (**Fig. 4**). The neurocranium had the lowest average thickness, whereas the dentary had the greatest thickness (**Fig. 4A**). The bathybenthic species *A. pallidus*, *A. thermalis*, and *A. platycephalus* consistently had the thickest bones. With the exception of the fifth ceratobranchial, *Cottocomephorus* and *Comephorus* had the thinnest ceratohyal, dentary, and neurocranium. As with mineral density, the reduction in thickness is convergent between both pelagic clades and is reflected across multiple bony elements.

**Fig. 4.**
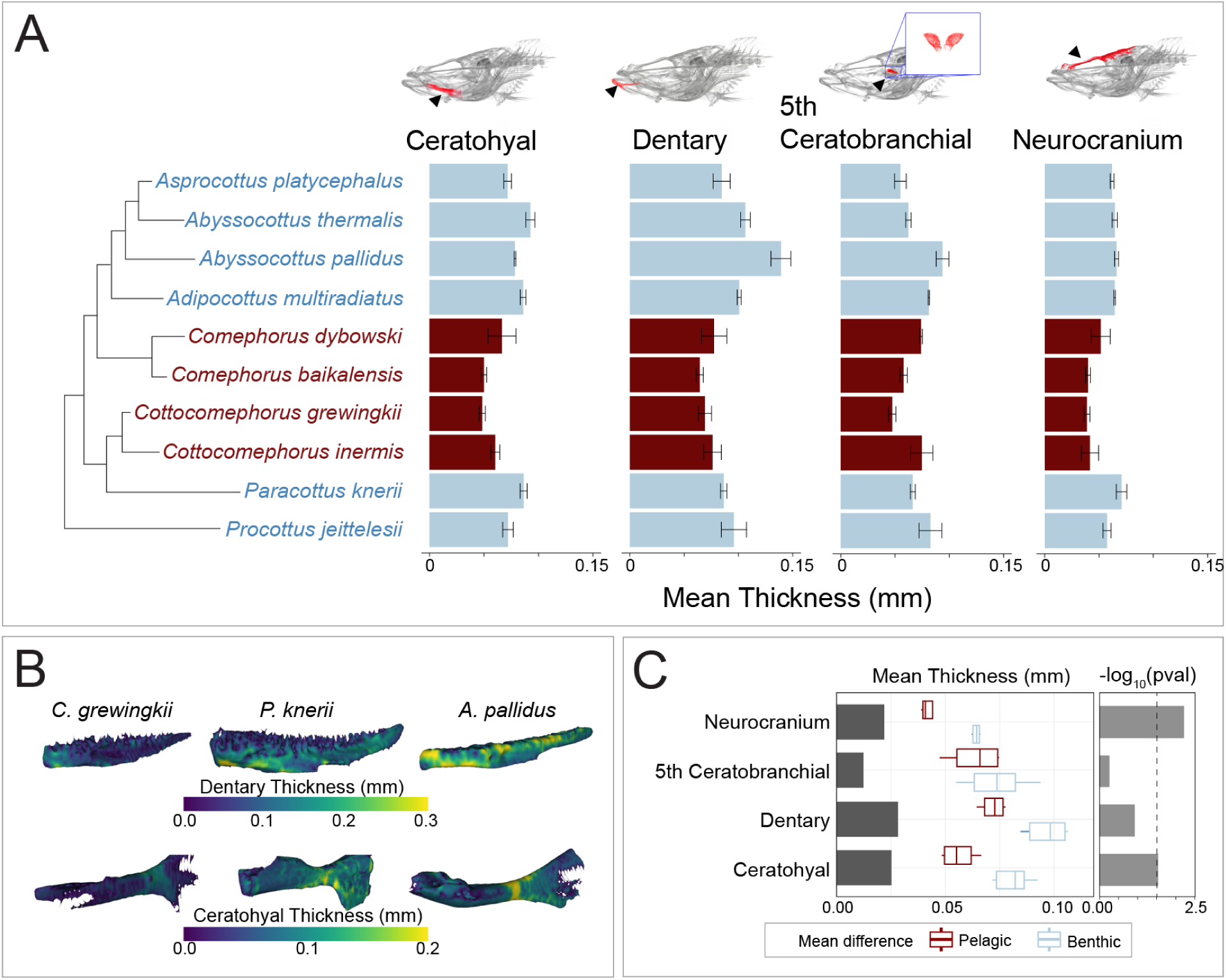
Variation in bone thickness across the Baikal sculpin phylogeny. A) Mean bone thickness in selected Baikal sculpin species, comparing benthic (blue) and pelagic (red) groups. B) Three-dimensional models of dentary and ceratohyal bones from representative specimens overlaid with a heatmap of bone thickness. C) Mean thickness differences between benthic and pelagic groups, with ANOVA *P*-values transformed via −log₁₀ (**Table S2**). Dashed line indicates a *P*-value of 0.05.

Across the three skeletal metrics of mineral density, porosity, and thickness, a consistent pattern emerged that separates pelagic from benthic Baikal sculpins. Pelagic species show lower mineral density, higher porosity, and thinner bones across multiple cranial elements, reflecting a repeated and multimodal shift toward lighter skeletal architecture. To evaluate whether these trait shifts align with ecological transitions, we examined how each bone metric varies with species position along the benthic to pelagic habitat axis using phylogenetic ANOVAs (**Table S2**). For mineral density (**Fig. 2C**, **Table S2**), water column position had significant effects in the basibranchial (*P* = 0.048), ceratohyal (*P* = 0.012), dentary (*P* = 0.004), and preopercle (*P* = 0.046), and marginal effects in the cleithrum (*P* = 0.074), fifth ceratobranchial (*P* = 0.062), and the neurocranium (*P* = 0.080). For bone thickness (**Fig. 4C**, **Table S2**), significant relationships with habitat were found for the neurocranium (*P* = 0.006) and the ceratohyal (*P* = 0.029), while no significant associations were detected for the dentary (*P* = 0.120) or the fifth ceratobranchial (*P* = 0.548). In contrast, bone porosity showed no statistically significant relationship with ecological position for any cranial element (**Fig. 3C**, **Table S2**). Taken together, the consistent trends toward lower mineral density, higher porosity, and reduced thickness in the bones of pelagic species, combined with element specific phylogenetic ANOVA results, indicate a repeated and convergent shift toward skeletal lightening in pelagic Baikal sculpins.

### Low correlations between bones and morphological variables

We next tested whether variation in density metrics was correlated across bones. To do this, we performed phylogenetic regressions using thickness, and porosity as predictors and mineral density as the response variable. These models showed very weak relationships between the three skeletal metrics, with adjusted r² values below 0.5 for all bones. Porosity showed a marginal association with mineral density only in the neurocranium (*P* = 0.0559), but model fit was still low (r² = 0.44), and thickness was not a significant predictor of mineral density for any bone examined. Separate regression models run for each predictor confirmed this result. Overall, there is little evidence that changes in mineral density, porosity, and thickness are tightly coordinated across cranial bones.

Because body size can influence bone density in fishes (Hamilton et al. 1981, St. John et al. 2022), we also examined correlations between centroid size and each skeletal metric. All bones exhibited weak negative correlations between body size and mineral density, porosity, and thickness (**Table 1**). This signal is driven largely by *Comephorus*, which had some of the largest specimens in the dataset. Overall, the influence of body size on cranial skeletal properties appears limited, and no bone showed strong evidence that larger individuals have systematically denser, less porous, or thicker cranial elements.

**Table 1.**
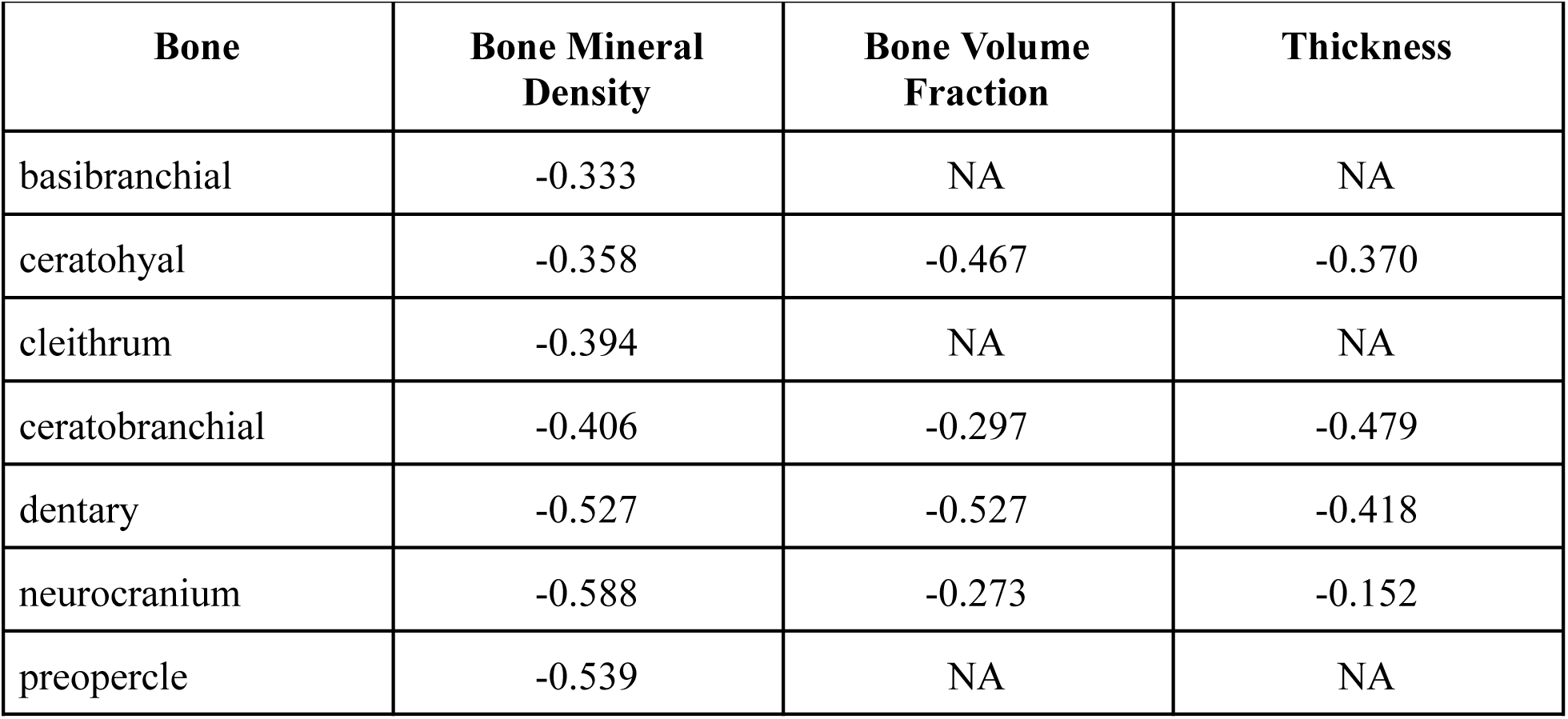
Spearman rank correlation coefficients between the various bone density and size metrics and centroid size.

### Convergent shifts in body shape during benthic-to-pelagic transitions

In addition to skeletal density, benthic-to-pelagic transitions typically involve coordinated changes to overall body shape, trending toward a slender, elongate body shape, furcate caudal fins, and a narrow caudal peduncle (Friedman et al., 2020; Rincon-Sandoval et al., 2020). Principal component analysis of shape data revealed distinct morphological patterns that are consistent with a recent analysis of Baikal sculpin shape variation (Sandel et al., 2025) (**Fig. 5A**). The first principal component (PC1) (54.31% of variation) clearly separated benthic and pelagic sculpins, while the second axis (19.92% of variation) primarily reflected differences among benthic species. Pelagic sculpins showed consistent morphological differences along PC1 relative to the benthic species, including smaller, posteriorly positioned eyes, wider fin bases, closer dorsal fin spacing, higher pectoral fin insertion, and longer, tapering tails (**Fig. 5B**). Although a multivariate phylogenetic ANOVA found no significant habitat effect on overall shape (F_(1,8)_ = 0.92, *P* = 0.44), a phylogenetic ANOVA focused on the PC1 scores revealed significant divergence between benthic and pelagic groups (F_(1,8)_ = 29.08, *P* = 0.02).

**Fig. 5.**
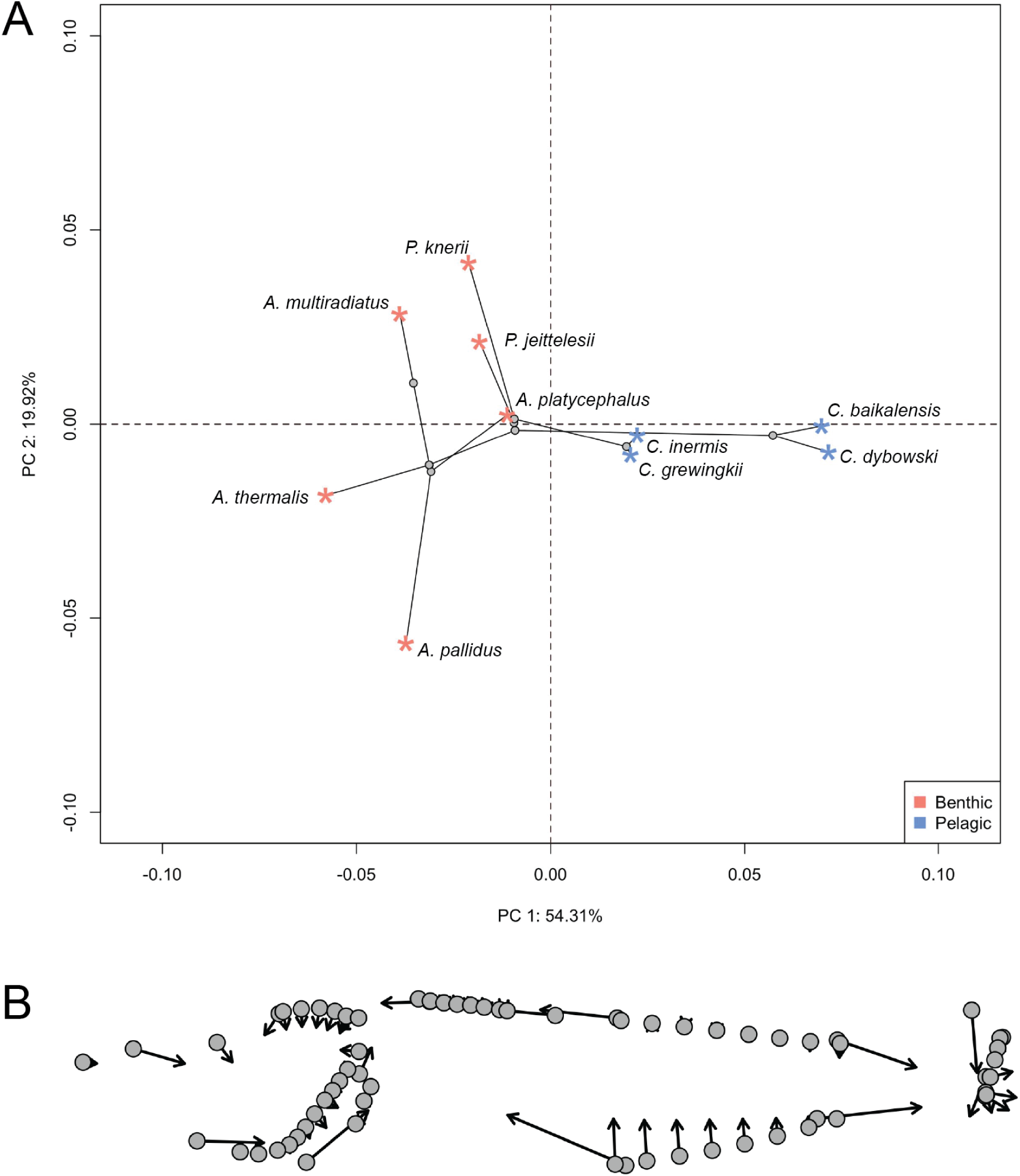
Convergent modifications to body shape during the benthic to pelagic transition in Lake Baikal sculpins. A) Phylomorphospace (i.e., a projection of the phylogenetic relationships in the morphospace) showing the first two axes of a principal components analysis of the shape data. B) Plot of vector displacements illustrating the changes in shape between the two extremes along PC1. Grey dots represent the extreme shape towards the negative end of the axis, while the tips of the vectors represent the extreme shape towards the positive end.

## Discussion

### Density patterns across the skull and phylogeny

Convergent reductions in skeletal density evolved in *Cottocomephorus* and *Comephorus* (**Figs. 2-4**), two genera that independently transitioned from benthic to midwater habitats (Sandel et al., 2025). The most conspicuous skeletal change in pelagic species was a skull-wide decrease in mineral density. However, a phylogenetic ANOVA revealed a significant association between mineral density and midwater habitat usage only in the dentary, ceratohyal, and basibranchial. Results in other bones are likely limited by small sample sizes, as our dataset includes only four pelagic species (out of five total), representing two evolutionary transitions (**Fig. 2**). Detecting significance in these elements, despite small sample sizes, implies that the divergence between mid-water and benthic phenotypes is substantial. Pelagic species also show changes in porosity and thickness, but these shifts are either inconsistent across pelagic taxa or are occasionally present in benthic species as well. *Comephorus baikalensis* had the lowest density across nearly every measurement, paralleling reports from Taliev (1955) and Sideleva (2003) of exceptionally low ash weight data and neutral buoyancy.

Density reductions affect skeletal elements differently, reflecting the unique functional demands of each bone. The basibranchials and neurocranium had the lowest density across metrics, with particularly low mineral density (**Fig. 2**), high porosity (**Fig. 3**), and reduced thickness (**Fig. 4**). Yet, aside from thickness, these bones did not exhibit the strongest habitat-specific density reductions (**Figs. 2-4**). By contrast, the dentary showed the largest mineral-density decrease between benthic and pelagic groups. Although associations between these density metrics and benthic and midwater habitats are not statistically significant after phylogenetic correction, there was consistently high porosity and low thickness within the pelagic jaws, especially the neutrally buoyant *C. baikalensis* (**Figs. 2-4**). Notwithstanding these reductions to density in pelagic species, the dentary was still denser than the other analyzed bones, a pattern also seen in low-density fishes such as snailfishes (Liparidae) and rattails (Macrouridae) (M. E. Gerringer et al., 2021; Martin et al., 2022). This combination of higher density feeding structures and a lower density neurocranium may reflect physical constraints on bone function: the dentary contends with high mechanical loading during feeding while the neurocranium primarily functions to protect the brain. Intriguingly, the primary predators of *Comephorus* are freshwater seals (*Pusa sibirica*), which lack typical molars used to crush prey and swallow these fishes whole (Valentina G. Sideleva, 2003; Yakupova et al., 2023). This predator-prey dynamic could influence selection pressures for either trait: relaxed selection for dense protective cranial bones in *Comephorus* and/or on molars in seals without hard-bodied prey. As mechanical load directly impacts fish bone density (Witten & Hall, 2015), the element-by-element variation across Baikal sculpins likely reflects a balance between genetic pathways favoring reduced density and plastic responses to mechanical load.

### Bone density: the depth axis

A defining ecological feature of Lake Baikal is its extreme depth, with basins exceeding 1,600 m (De Batist et al., 2006). The swim bladder, the primary buoyancy organ in most fishes, becomes increasingly ineffective with depth, as hydrostatic pressure both impedes inflation and compresses gas to high densities (Marshall, 1960; Priede, Jamieson, Bond, & Kitazato, 2024). Although some deepwater marine species with swim bladders occur even at hadal depths, such as the cusk eel *Bassogigas profundissimus* (7,160 m) and the rattail *Coryphaenoides yaquinae* (7,259 m) (Nielsen & Munk, 1964; Priede et al., 2024), many deep-sea fishes lack one. Instead, several lineages rely on low-density lipids to maintain buoyancy, as in deep-sea bristlemouths and diel-migrating lanternfishes (Blaxter, Wardle, & Roberts, 1971; Crabtree, 1995; Godø, Patel, & Pedersen, 2009; Neighbors & Nafpaktitis, 1982; Priede, 2017). Additional weight-reducing modifications occur in soft tissues: some species possess low-density “watery” muscles (Crabtree, 1995; Priede, 2017; Tuponogov, Orlov, & Kodolov, 2008), gelatinous subdermal matrices with moderately positive buoyancy (Mackenzie E. Gerringer et al., 2017), or enlarged, fluid-filled crania (Horn, Grimes, Phleger, & McClanahan, 1978).

Bone, one of the densest tissues, often shows reduced density across deep-sea fishes, leading to the hypothesis that skeletal density decreases with depth in fishes (Childress & Nygaard, 1973; Cohen, 1974; Denton & Marshall, 1958; M. E. Gerringer et al., 2021). Baikal sculpins show no apparent association between depth and bone density, as the bathybenthic clade has densities similar to shallow benthic species (**Figs. 2-4**). This pattern mirrors that of rattails, which also show no depth-density correlation, though rattails do possess swim bladders (Martin et al., 2022). Instead, bone density in sculpins is correlated with benthic and midwater habitat usage, which was also observed in snailfishes; pelagic snailfishes display lower skeletal density, fewer vertebrae, and loss of bony elements such as suction disks (M. E. Gerringer et al., 2021). Thus, correlations between skeletal density and depth appear to reflect midwater habitat usage and the presence or absence of a swim bladder rather than an inherent product of depth adaptation.

### Contrasts to the cryonotothenioid adaptive radiation

Skeletal reduction patterns in Baikal sculpins closely parallel those in Antarctic notothenioids, though the changes to skeletal density within the two clades differ in evolutionary timing and mode. Skeletal density was initially reduced prior to the cryonotothenioid adaptive radiation, with the skeleton becoming even less dense in secondarily pelagic groups (Daane et al., 2019; Eastman et al., 2014). In contrast, Baikal sculpins appear to have evolved reduced density independently within the secondarily pelagic lineages without being preceded by changes to density in benthic relatives (**Figs. 2-4**).

Our dataset does not allow direct verification of cartilage retention, but the low densities we observed are consistent with highly cartilaginous skeletons, as in notothenioids (Eastman, 2024). Interestingly, comparisons of benthic notothenioids to benthopelagic icefishes (Channichthyidae) revealed no consistent differences in mineral density or porosity among bones despite variation in overall density, leading the authors to suggest that bone shape and changes in developmental timing, known as heterochrony, may contribute more strongly to density reduction than bone mineral density and microstructure (Ashique et al., 2022). Many benthic sculpins and notothenioids are pelagic as larvae or juveniles (T. J. Buser, Finnegan, Summers, & Kolmann, 2019; Eastman, 2024). Retention of juvenile characters into adulthood could facilitate midwater adaptation in these clades (Eastman, 2024; Eastman et al., 2014; Taliev, 1955). A similar pattern is also observed in pelagic snailfishes, which retain larval-like morphologies into adulthood (Marliave & Peden, 1989). Exploration of potential patterns of paedomorphy in Baikal sculpins should be a focus of future investigations into their skeletal evolution.

### Pelagic body shape: evolutionary parallelism

Beyond reduced skeletal density, we observe convergent shifts toward elongated body shapes in the pelagic sculpins (**Fig. 5**). Elongate morphologies reduce drag and enhance swimming efficiency, whereas deep-bodied benthic forms favor maneuverability within structured habitats but at the cost of swimming speed (Hatfield & Schluter, 1999; P. Webb, 1984; P. W. Webb, 1982). Similar transitions have evolved repeatedly across teleosts during benthic-to-pelagic shifts, in species both with and without swim bladders (Cooper et al., 2010; Friedman et al., 2020; Rincon-Sandoval et al., 2020; Robinson & Wilson, 1994; Walker, 1997). The mechanistic basis of this convergence remains unresolved but likely reflects shifts toward a hydrodynamic optimum for swimming efficiency, as well as developmental and genetic constraints that could channel evolution along similar mutational trajectories during benthic to pelagic transitions.

Notably, body elongation is also observed in *Leocottus*, the benthic sister group to *Cottocomephorus*, though to a lesser degree than in the pelagic species (St. John et al., 2022). *Leocottus* further clusters with *Cottocomephorus* in morphospace (Sandel et al., 2025), suggesting similar morphologies may have arisen prior to the benthic-to-pelagic transition of *Cottocomephorus*.

We also find evidence for convergent evolution in fin placement between benthic and pelagic sculpins. While benthic sculpins possess lower, ventrally inserted pectoral fins used for station-holding on the substrate, pelagic species display pectoral fins inserted higher on the flanks. This higher, more horizontal orientation is thought to help the benthopelagic *Cottocomephorus* transition more readily from resting to forward propulsion (Taliev, 1955). Although not quantified here, pectoral fins are also greatly elongated in *Comephorus* and *Cottocomephorus*, which increases surface area for stabilizing hydrodynamic drag and buoyancy (Jakubowski, Tugarina, & Zuwała, 2003; Valentina G. Sideleva, 2003). This functional trade-off parallels the silverspotted sculpin, *Blepsias cirrhosus*, where reduced ventral fin development and higher aspect ratios correlate with epi-benthic sculling rather than anchoring (Kane & Higham, 2012). Thus, shifts in fin insertion likely reflect a dual pressure: the loss of station-holding utility and the increased demands of pelagic hydrodynamics.

Cottid fishes are almost entirely benthic with highly conserved morphologies (Thaddaeus J. Buser et al., 2024; T. J. Buser et al., 2019). The divergence from benthic morphospace and reduction in skeletal density in pelagic Baikal sculpins highlights an ancestral capacity within cottids to diversify along the benthic-pelagic axis. This potential is likely constrained in their shallow freshwater relatives by a lack of ecological opportunity, specifically the shallow nature of most lakes and streams and lower prey diversity in boreal and temperate climates (Thaddaeus J. Buser et al., 2024). Notably, only five of the ∼40 Baikal species are pelagic or benthopelagic (Sandel et al., 2025), a pattern also consistent with a broader trend in teleost fishes in which benthic clades are typically more speciose and morphologically diverse than pelagic ones (Friedman et al., 2020; Rincon-Sandoval et al., 2020).

### Pathways to pelagicism

The genetic and developmental mechanisms underlying these skeletal density reductions and body shape changes remain unresolved. Convergent evolution within Baikal sculpins, alongside broader parallels across Perciformes, including snailfishes (M. E. Gerringer et al., 2021) and notothenioids (Eastman et al., 2014), positions this clade of fishes as a powerful comparative system for gene discovery. The repeated evolution of similar skeletal phenotypes across pelagic perciform taxa provides a foundation for the use of comparative genomic tools to identify molecular pathways undergoing parallel evolutionary changes in tandem with convergently evolved traits (Hilgers & Hiller, 2025; Hiller et al., 2012).

A potential cellular mechanism for skeletal density reduction in sculpins and notothenioids involves the shared developmental origin of osteoblasts and adipocytes. Both secondarily pelagic sculpins and notothenioids are exceptionally lipid-rich in multiple tissues. For example, *Comephorus baikalensis* accumulates triglycerides during early growth, eventually reaching a remarkable ∼39% lipid by wet weight (Kozlova & Khotimchenko, 2000). Lipid accumulation also occurs in the neutrally buoyant notothenioids such as *Pleuragramma antarcticum*, which possesses specialized subcutaneous and intramuscular lipid sacs (Devries & Eastman, 1978; Eastman, 1988). Further, other deepwater marine fishes exhibit lipid-rich bone tissue (Lee, Phleger, & Horn, 1975), and, in mammals, osteoporosis is associated with elevated bone marrow adiposity (Muruganandan, Govindarajan, & Sinal, 2018). Because osteoblasts and adipocytes arise from mesenchymal stem cells (Aghebati-Maleki et al., 2019; Chen et al., 2016; Hu et al., 2018; Pittenger et al., 2019), these observations suggest possible correlated trait evolution between skeletal density reduction and increased lipid storage via regulation of mesenchymal stem cell differentiation. Comparative genomic tools may be useful in unraveling the molecular mechanisms underlying these convergently evolving, correlated changes to lipid metabolism and skeletal density.

### Summary

Here, we characterized patterns of skeletal density across the radiation of Baikal sculpins. We found that pelagic lineages have independently evolved reduced skeletal density, likely as a buoyancy mechanism. We found a multi-modal reduction in skeletal density within midwater fishes based both on mineral content and microstructure, though the contributions of each mode of density reduction depended on the specific bones and species. These reductions in skeletal density were accompanied by convergent overall changes to body shape, including more streamlined morphologies and a shift in the position of the pectoral fins, mirroring a common teleost-wide trend in benthic-to-pelagic transitions. Future work should leverage these convergent phylogenetic patterns to dissect the molecular basis of skeletal density evolution and benthic-to-pelagic transitions.

## Acknowledgements

This work was supported by the US National Science Foundation (NSF) grants OPP-2324998 to J.M.D. and DEB-1557147 to M.W.S and A.A., a US National Institutes of Health (NIH) grant R35GM150590 to J.M.D, and the Limnological Institute SB RAS 1025032400008-1-1.5.12 to S.K.

## Author contributions

B.A.G., O.L., J.M.D. designed research; M.W.S., A.A. and S.K. collected and provided key samples; B.A.G., O.L., S.L., M.P.N., K.M.E. performed research; B.A.G., O.L., M.E.G. and J.M.D. analyzed data; and B.A.G., O.L. and J.M.D. wrote the paper; B.A.G., O.L., S.L., M.P.N., M.E.G., K.M.E., A.A., S.K., M.W.S. and J.M.D. provided critical revision of the manuscript.

## Supplemental Methods

### Micro-CT Scan and Reconstruction Settings

Specimens were scanned using a Bruker SkyScan 1273 micro-CT scanner configured with 70 kV X-ray voltage, 214 µA current, medium focal spot size, and 0.3° rotation step. Scan lengths ranged from 145 to 245 mm with voxel sizes between 12 and 20 µm, optimized for each specimen’s size. Two calcium hydroxyapatite phantom markers (25% and 75% density) were included in each scan for bone mineral density calibration.

Micro-CT data reconstruction used Bruker NRecon v.2.1.0.1, cropping the region from the snout to the 6th or 7th vertebra. Reconstruction involved histogram adjustments with a zero lower bound and standardized upper bound. Data downsampling in DataViewer employed “Single Volume of Interest” with resize factors of 2 or 3, no color or intensity corrections were performed during downsampling.. Amira segmentation created LabelFields using paintbrush tools with interpolation (Ctrl+I), while phantoms were segmented separately using masking functions and magic wand tools with consistent pixel brightness thresholds. Ring artifact reduction set to six for improved image quality. For oversized specimens, we performed multiple overlapping scans stitched into composite datasets. Large files were reduced using Bruker DataViewer v.1.5.4 with downsampling factors of 2 or 3. This standardized protocol with phantoms in every scan ensured consistency across specimens and species, making bone property data directly comparable among Baikal sculpins in the present dataset.

### Computation of Bone Mineral Density

Material statistics were extracted from Amira’s measurement tool, which provided voxel count and cumulative brightness sum for each segmented region. Mean pixel brightness (MPB) was then computed separately as:

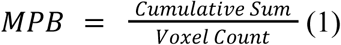

To establish a reference model, the MPB values of the 25% and 75% hydroxyapatite phantoms were used to create a linear regression model, correlating MPB with known density percentages. The general form of this model was:

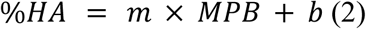

where m represents the slope of the calibration curve derived from the phantom MPB values and their known hydroxyapatite percentages, and b is the intercept.

Once the calibration model was established, the MPB of each segmented bone was calculated from the material statistics and substituted into the model to determine its hydroxyapatite percentage:

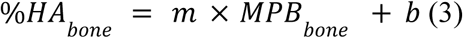

This method ensures that bone mineral density quantification remains standardized across different scans and specimens.

### Standardized Density Visualization

To standardize visualization across scans, colormap thresholds were adjusted to represent values from 10% to 90% hydroxyapatite:

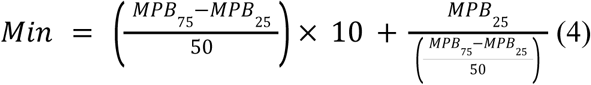

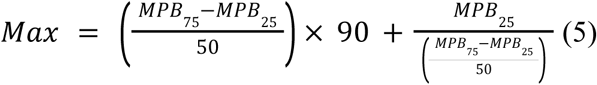

These thresholds ensured that pixel intensities outside the target range were rendered as black (low) or white (high) for visual contrast.

### Image Stack Loading and Visualization

We imported reconstructed scan data consisting of log files and PNG slices into 3D Slicer using the Image Stacks module. The pixel size extracted from log files was converted from micrometers to millimeters and entered into the Spacing field in the Volumes Module under Volume Information. By adjusting slice skip parameters, we maintained total output sizes below 200 MB to ensure computational efficiency. For optimal bone visualization, we enabled volume rendering using the CT AAA preset while making adjustments to opacity mapping and region of interest boundaries.

### Segmentation and Isolation of Bone Regions

Using the Segment Editor module, we performed segmentation by first applying consistent thresholding across specimens (min: 30, max: 255) to standardize pixel intensity selection. We then refined the segmentations using paint, erase, and scissors tools to isolate bones of interest from surrounding structures. These segmented regions represented the bone volume (BV) used in subsequent bone volume fraction calculations.

### Generation of Total Volume (TV)

We created solid bone representations by cloning each isolated segmentation in the Data module and applying the Wrap Solidify tool to generate watertight total volumes (TV). This process filled both internal and external voids. When necessary, we performed additional manual segmentation refinements using paint, erase, and fill between slices tools.

### Bone Volume Fraction Calculation

We calculated bone volume fraction (BVF) as the ratio of bone volume (BV) to total volume (TV). Volume measurements were obtained through the Segment Statistics module, extracting BV and TV values using Label Map Statistics, Scalar Volume Statistics, or Closed Surface Statistics. Valid BVF values ranged between 0 and 1, with values exceeding 1 indicating segmentation errors that required corrections to the total volume segmentation.

### Bone Thickness Measurement

For thickness measurements, we exported BV segmentations as both Labelmap and Model files. Using the Simple Filters module, we processed these files through a binary thinning algorithm (BinaryThinningImageFilter) followed by a distance transformation (DanielssonDistanceMapImageFilter) to compute thickness at each model point. The Probe Volume With Model module then generated 3D thickness heatmaps where yellow indicated thicker regions and dark blue represented thinner areas.

### Computation of Mean Bone Thickness

We computed mean bone thickness using Python scripting within 3D Slicer. The script processed segmented bone models by extracting thickness values from thousands of surface points. For each model, the script retrieved thickness values stored in the point data and calculated the mean by averaging all measurements. This semi-automated approach ensured consistent specimen comparisons and enabled efficient quantification of mean bone thickness and porosity. The resulting standardized metrics revealed structural patterns that complemented our bone mineral density assessments.

### Standard Visualization

Micro-CT scan images of our Baikal sculpins (Figure 7) were processed in Amira using colormap settings calibrated to the minimum (4) and maximum (5) values established in Section 2.5.5. These threshold values, representing the 10% to 90% hydroxyapatite range, were selected to exclude potential measurement uncertainties at the extremes of our data range.

## Supplemental Figures

**Fig. S1.**
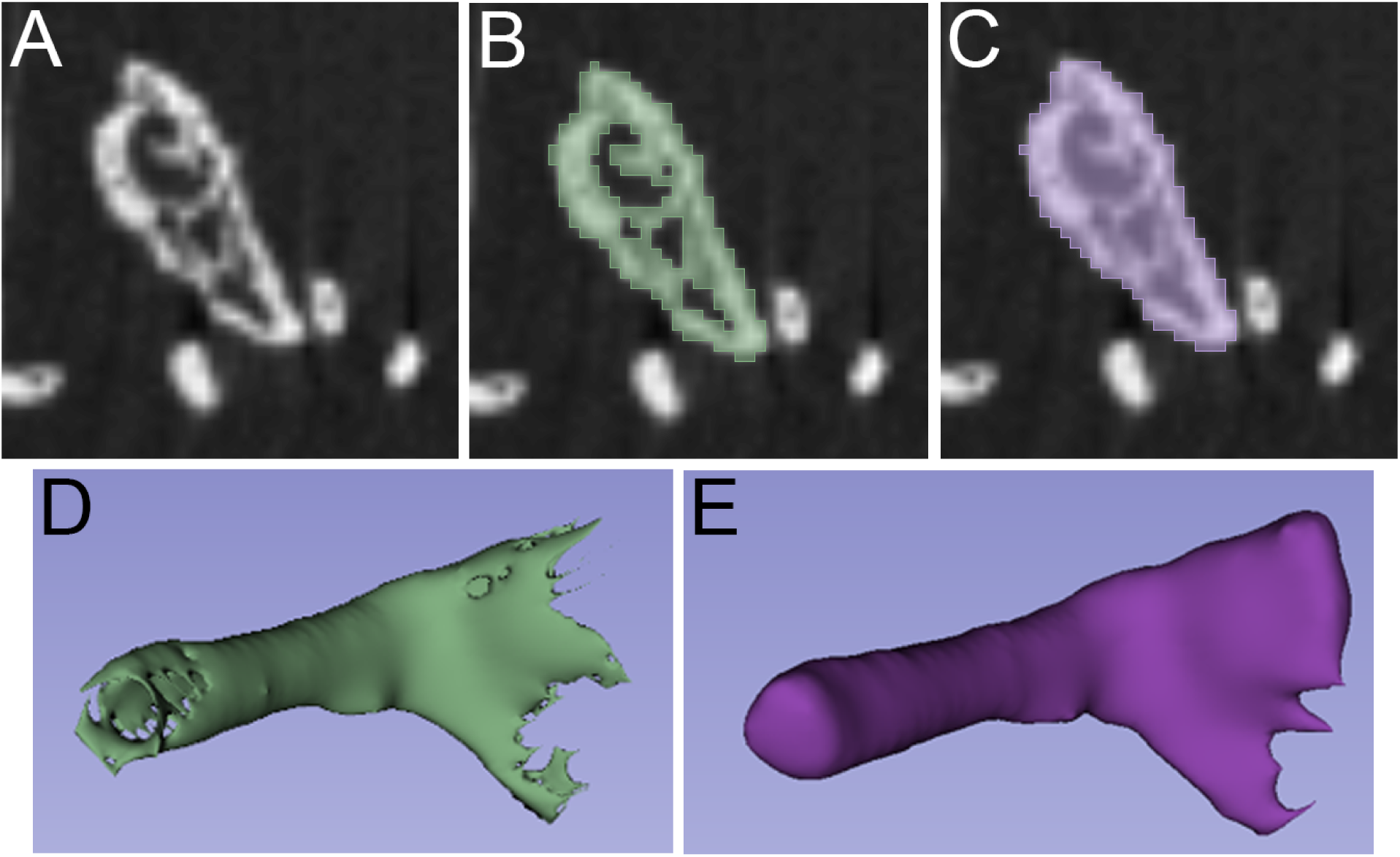
Methodology for estimating bone volume fraction (BV/TV). A) Representative CT cross-section of the ceratohyal. B) Segmentation of mineralized tissue used to calculate Bone Volume (BV). C) Total Volume (TV) defined by filling internal voids. D) 3D reconstruction of the segmented BV. E) 3D reconstruction of the TV.

**Fig. S2.**
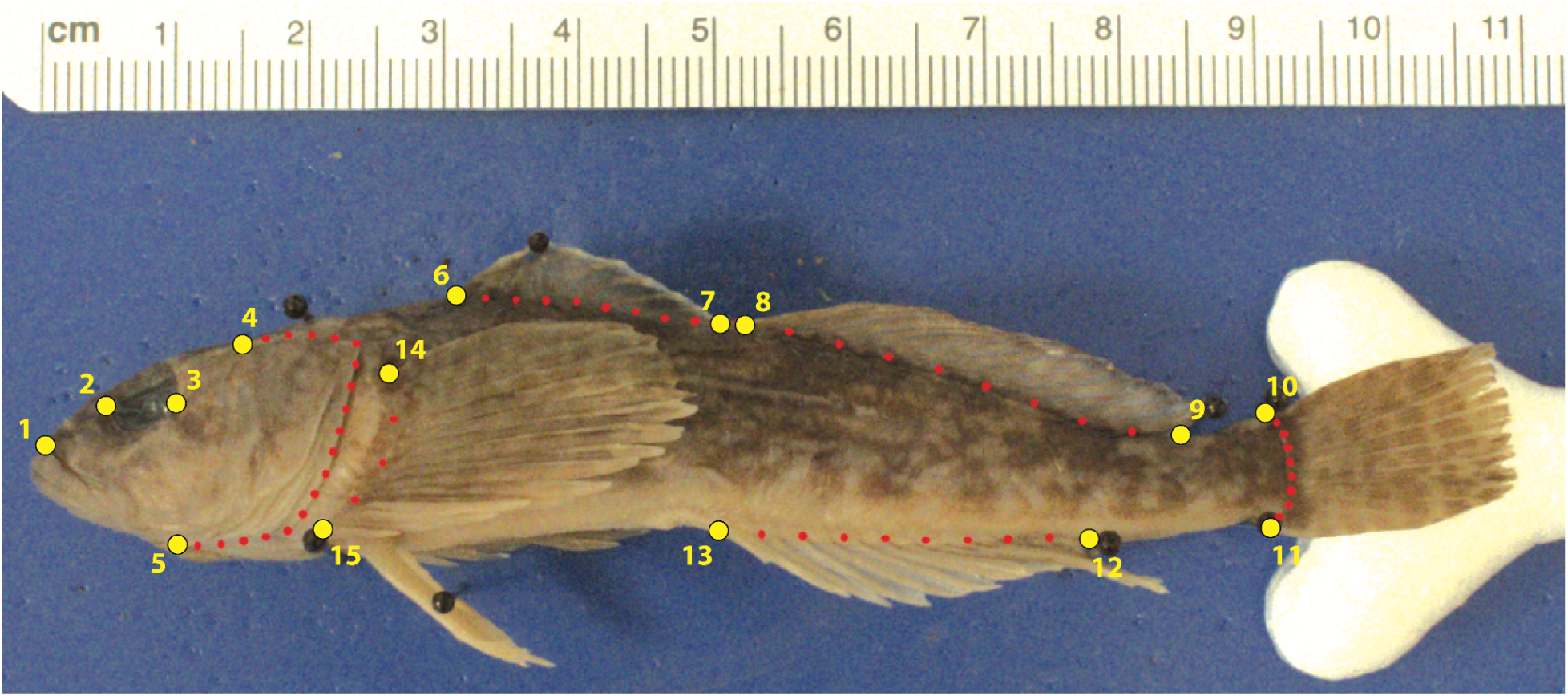
Anatomical landmarks for geometric morphometrics. Example with *Paracottus knerii*. Primary landmarks (yellow): snout tip (1), orbital width (2-3), operculum (4-5), base of spinous dorsal fin (6-7), base of soft-rayed dorsal fin (8-9), base of caudal fin (10-11), base of anal fin rays (12-13), base of pectoral fin (14-15). The semi-landmarks are listed in red. There were 18 semi-landmarks for the operculum and 8 semi-landmarks each for the base of the spinous and soft-rayed dorsal fins, the caudal fin, and the anal fin. There were 3 semi-landmarks used for the base of the pectoral fin.

## Supplemental Tables

**Table S1.** List of anatomical landmarks and semi-landmarks used for geometric morphometrics.

| **Anatomical Structure** | **Anchor Landmarks Number(s)** | **Number of Semi-Landmarks** |
| --- | --- | --- |
| Snout Tip | 1 | n/a |
| Orbital Width | 2,3 | n/a |
| Operculum | 4,5 | 18 |
| Base of spinous dorsal fin | 6,7 | 8 |
| Base of soft-rayed dorsal fin | 8,9 | 8 |
| Base of caudal fin | 10,11 | 8 |
| Base of anal fin Rays | 12,13 | 8 |
| Base of pectoral fin | 14,15 | 3 |

**Table S2.** Average phylogenetic ANOVA *P*-values for difference in bone properties between benthic and pelagic species. n.t. indicates not tested.

| **Bone** | **Bone Mineral Density** | **Bone Volume Fraction** | **Thickness** |
| --- | --- | --- | --- |
| basibranchial | 0.04773 | n.t. | n.t. |
| ceratohyal | 0.01185 | 0.49330 | 0.02910 |
| cleithrum | 0.07463 | n.t. | n.t. |
| ceratobranchial | 0.06247 | 0.83746 | 0.54766 |
| dentary | 0.00422 | 0.16758 | 0.11963 |
| neurocranium | 0.08007 | 0.14794 | 0.00607 |
| preopercle | 0.04607 | n.t. | n.t. |

